# Exogenous stimulation of *Tanacetum vulgare* roots with pipecolic acid leads to tissue-specific responses in terpenoid composition

**DOI:** 10.1101/2024.04.28.591506

**Authors:** Humay Rahimova, Robin Heinen, Baris Weber, Wolfgang W. Weisser, Jörg-Peter Schnitzler

## Abstract

*Tanacetum vulgare* L., known as tansy, is a perennial plant with a highly variable terpenoid composition, with mono- and sesquiterpenoids being the most abundant. The high diversity of terpenoids is known to play an important role in mediating ecological interactions. However, the distribution of terpenoids in different tissues and the inducibility of terpenoids in these tissues by biotic stress are poorly understood.

In this study, we investigated the changes in terpenoid profiles and concentrations in different plant organs following treatment of roots with pipecolic acid (Pip). Pipecolic acid is a non-proteinogenic amino acid that triggers defense responses in plants. It is often used to induce systemic resistance (SAR) in plants under controlled conditions.

Examination of the tissues showed that the leaves and midribs contained mainly monoterpenoids, while the coarse and fine roots of the plants contained mainly sesquiterpenoids. The rhizomes occupied an intermediate position by presenting the terpenoid profiles of both the midribs and roots but also the unique compounds of its own. Treatment with pipecolic acid led to an increase in the concentration of mono- and sesquiterpenoids in all tissues except rhizomes. However, a significantly higher amount of sesquiterpenoids was formed in root tissues in response to Pip compared to shoots.

The metabolic atlas for terpenoids presented here shows that there is an exceptionally strong differentiation of terpenoid patterns and terpenoid contents in the different tissues of tansy. This, together with the differential inducibility by biotic stress, suggests that the chemical diversity of terpenoids may play an important role in the ecological interactions of tansy and in the defense against biotic stressors that feed on the below- and above-ground organs of the plant.

## Introduction

Specialized metabolites are a diverse group of compounds produced by plants that play a crucial role regulating interactions with the environment. The production of specialized metabolites in plants is regulated by transcriptional mechanisms, resulting in the tissue-specific synthesis of compounds - such alkaloids and terpenoids - that play a role in mediating interactions with the biotic and abiotic environment (Erb & Reymond, 2019; Wetzel & Whitehead, 2020). Previous studies on metabolic variation in response to abiotic and biotic factors have mostly examined metabolic responses in plant shoot tissues, but, given some of the practical challenges, comparatively little is known about how different abiotic or biotic factors alter the metabolic profiles of different types of belowground tissues (Rasmann et al., 2012; van Dam, 2009). Considering that shoots and roots are exposed to divergent herbivore and pathogen communities, they may face unique selection pressures. The production and distribution of secondary metabolites such as flavonoids, alkaloids, and terpenoids in plants may vary between different plant organs like shoots and roots (Kleine & Müller, 2013; Zhang et al., 2021). However, the metabolic profile of structures and organs at finer scales is still largely unknown.

*Tanacetum vulgare* L. (Asteraceae species), commonly referred to as tansy, is a fragrant herb native to Eurasia. Its terpenoid content varies considerably across individuals, as evidenced by studies conducted by Clancy et al. (2016), Kleine & Müller (2011), and Rahimova et al. (2024). Tansy leaves contain glandular trichomes on the tissue surface that store the terpenoids. These terpenoids can be induced and emitted immediately upon insect and herbivory attacks (Clancy et al., 2016, 2020). Tansy terpenoids exhibit high intraspecific variability, allowing for the classification of plants into distinct chemotypes based on their dominant or mixed compounds, in the absence of a discernible dominant compound (Dussarrat et al., 2023; Rahimova et al., 2024). The diverse group of terpenoids in tansy has been demonstrated to influence plant-insect interactions (Ojeda-Prieto et al., 2024). Terpenoids have been found to correlate with plant-associated aphid communities, and they can also determine aphid feeding preference in tansy leaves (Clancy et al., 2016; Jakobs and Müller, 2018; Neuhaus-Harr et al., 2024). However, to date, there is no information available about the behavior of terpenoid profiles upon induction by stressors, and whether this varies between tissues, such as leaves, rhizomes, and root tissues.

Terpenoids represent the largest group of specialized (secondary) metabolites in plants and play an important role in various biological processes, especially in the interactions of plants with their environment and other organisms (Pichersky & Raguso, 2018). This extensive variety of terpenoids is regulated by terpene synthases, which are the most significant enzymes involved in the synthesis of hemiterpenes (C5), monoterpenes (C10), sesquiterpenes (C15) or diterpenes (C20) (Bohlmann et al., 1998; Tholl, 2006). Of these subgroups, the mono- and sesquiterpenes represent the predominant group in tansy and can be localized in different tissues. For instance, the shoot tissues are primarily composed of the C10 compound group, the monoterpenes (Kleine & Müller, 2013; Clancy et al., 2016), whereas the C15 group, the sesquiterpenes, are predominantly found in roots (Kleine & Müller, 2013). Terpenoids can act as direct defenses against herbivores and pathogens, influencing the interactions between plants and their environment (Penuelas et al., 2014). Due to their volatility, they also serve as indirect defenses (‘cry for help’), attracting the predators of herbivores that attack them (Dicke and Baldwin, 2010). For instance, switchgrass (*Panicum virgatum* L.) plants emit a combination of mono- and sesquiterpenes when attacked by the generalist herbivore *Spodoptera frugiperda* (J.E. Smith) (fall armyworm) and when there is a systematic response to root treatment with defense hormones such as methyl jasmonate (Muchlinski et al., 2019). Some of these compounds, especially monoterpenes, play a role not only in defense against pathogens and insects, but also in adaptation to environmental challenges such as drought and high temperatures (Vaughan et al., 2015; Zuo et al., 2017). Other terpenoids, such as the sesquiterpene β-caryophyllene, also found in tansy (Rahimova et al., 2024), influence heliotropism and control leaf movement (Nilsen & Foresth, 2018) and root growth and hydrotropism (Yamagiwa et al., 2011). Induced by herbivory on maize roots, it acts as an attractant for entomopathogenic nematodes (EPNs) (Turlings et al., 2012).

Plants are constantly under attack from a variety of biotic factors such as insects, herbivores, viruses, fungi and bacteria, which can affect plant health and productivity (Choudhary & Senthil-Kumar, 2022; Irulappan et al., 2022; Poelman & Kessler, 2016). When a plant is challenged by biotic attackers, a number of biochemical reactions occur in the locally infected plant tissues that can lead to systemic acquired resistance (SAR) (Vlot et al., 2021) or induced systemic resistance (ISR). While the latter is mediated by the jasmonic acid (JA)/ethylene pathway, the induction of SAR is dependent on salicylic acid (SA) (Vlot et al., 2009) Aphid feeding stimulates SA production (Voelckel et al. 2004), whereas leaf-chewing herbivores induce the JA/ethylene pathway (Walling 2000). JA-induced defense mechanisms have been shown to impair aphid performance, whereas SA-induced defense mechanisms have less negative effects (Agrawal 1998; Ali & Agrawal 2014). Pipecolic acid (Pip) is a positive regulator of SA production and leads to an increase in the plant’s defenses (Chen et al., 2018; Hartmann et al., 2018; Vlot et al., 2021). For example, when *Arabidopsis thaliana* (L.) Heynh. plants are exposed to microbial infection, Pip, a non-protein amino acid, is synthesized from L-lysine (Návarová et al., 2013) and then undergoes N-hydroxylation to produce N-hydroxypipecolic acid (NHP) (Hartmann et al., 2018). Both Pip and NHP are mediators of SAR and accumulate in locally infected and distal tissues during infection (Hartmann et al., 2018). Furthermore, Pip/NHP triggers the production of volatile organic compounds (VOCs) in plants as part of the SAR mechanism, contributing to intra- and inter-plant defense propagation through the emission of VOCs that act as defense cues (Brambilla et al., 2023; Vlot et al., 2021). When SAR is induced by bacterial infection in *Arabidopsis*, plants emit VOCs containing elevated levels of monoterpenoids such as α-pinene, β-pinene and camphene that triggers plant to plant communication (Wenig et al., 2019) within and between species (Frank et al., 2021). In barley (*Hordeum vulgare* L.), Pip drip irrigation similarly induces defense responses against subsequent powdery mildew infection (Lenk et al., 2019). Given that Pip application to roots provides a robust mimic of biotic induction in many model systems, and that the Pip pathway is widely reported across the plant kingdom, this provides an excellent opportunity to use Pip to study differences in terpenoid inducibility between different plant organs of tansy.

In this study, we mimicked the SA-dependent response of tansy to aphid attack leading to SAR by using Pip as a stimulant and investigated the effects of exogenously applied Pip on tansy defense responses. We ask:

1. How does the terpenoid content of tansy differ between five major plant tissues: coarse roots, fine roots, rhizome, leaflets and midribs? Overall, we aimed to compile a metabolome atlas on the occurrence of terpenoids and their chemical diversity in tansy. We hypothesized that there are tissue-specific differences in the overall terpenoid atlas of tansy, as different terpenoid group distributions between shoots and roots have been reported in the literature. Next, we asked:
2. Does Pip induce terpenoids as a defense response equally in tansy shoot and root tissues? We predicted that Pip will increase the induction of terpenoids mainly in root tissues, but also in shoot tissues. We expected a stronger response of sesquiterpenoids in belowground and monoterpenoids in aboveground plant organs.

## Material and Methods

### Tansy cultivation and pipecolic acid treatment

The tansy plants used in this study originated from a stock of 120 plants propagated from seeds collected from twelve individual mother plants on plots of “The Jena Experiment” (https://the-jena-experiment.de), Jena, Germany in 2020. Five unique genotypes were collected for further propagation. Plants were clonally propagated by rhizome cutting in a greenhouse (21.0 °C, 14:10 h light:dark) in February 2022. After successful root formation, the plantlets were transplanted into pots (10 x 10 x 11 cm) with commercial soil (Floradur Bodensubstrat, Floragard Vertriebs-GmbH, Oldenburg, Germany) and fertilized with Hakaphos® Red (8% N, 12% P2O5, 24% K2O, 4% MgO, 31% SO3, 0. 01% B, 0.02% Cu, 0.05% Fe, 0.05% Mn, 0.001% Mo, 0.02% Zn; Compo Expert GmbH, Münster, Germany). Nine weeks after propagation, twenty of the resulting forty plants were randomly assigned to a Pipecolic acid (Pip) treatment and twenty to a control group, irrespective of the genetic identity of the plants. Forty mL of Pip solution (10 µM Pip in H_2_Obidest, Sigma Aldrich, Taufkirchen, Germany) was applied to each pot by watering the root system. The control group received the same treatment but without the addition of Pip. Three days later, on April 7^th^, and 8^th^ 2022, all plants were harvested. At harvest, the tansy plants were divided into five different tissues: (i) Leaflets, (ii) leaf midrib, (iii) rhizomes, (iv) coarse roots and (v) fine roots. To avoid induction by mechanical damage, samples were processed within one minute (each fresh tissue was cut and packaged separately) and frozen in liquid nitrogen to stabilize molecular processes.

### Hexane extraction of terpenoids and GC-MS analyses

Chemical analyses were performed as reported by Clancy et al. (2016), with minor modifications. Frozen samples were ground to a fine powder and immediately stored at −80°C until further processing. Six hundred µL of hexane containing 860 pmoL µL^−1^ of internal standard (monoterpene δ-2-carene) was added to 300 mg of frozen samples, and the mixture was vortexed and refrigerated at 4°C for 24 hours. Then 150 µL of the extract was removed and stored at 4°C. A further 150 µL of n-hexane was added a second time, vortexed and stored for 24 hours. One hunded and fifty µL of hexane extract was again collected and combined with the liquid extract initially collected. The samples were analyzed in a gas chromatography-mass spectrometer (GC-MS) by injecting 1 µL into a glass microvial in an empty glass cartridge placed on an autosampler. The samples were desorbed from 35°C to 240°C at a rate of 120°C min^−1^ (held for 2 min) in the thermal desorption unit (TDU, Gerstel, Mülheim an der Ruhr, Germany) connected to a GC-MS (GC category: Agilent Technologies, 7890A, MS category: 5975C inert XL MSD with a triple axis detector, Palo Alto, CA, USA) using an arylene siloxane capillary column (60 m × 250 μm × 0.25 μm DB-5MS + 10 m DG, Agilent Technologies) with a mixture of 5% phenyl 95% dimethyl. Samples were analyzed in splitless mode at a constant flow rate of carrier gas (He: 1 mL min^−1^). The temperature program was 40°C (held for 0 min) to 150°C with a ramp rate of 10°C min^−1^, then 80°C min^−1^ to 175°C, then 5°C min^−1^ to 190°C, then 80°C min^−1^ to 250°C, then 100°C min^−1^ to 300°C and held for 6 min. The chromatograms from the GC-MS analysis were evaluated, integrated and quantified using Enhanced ChemStation software (MSD ChemStation E.02.01.1177, 1989-2010 Agilent Technologies, Santa Clara, CA, USA). The peaks were first checked for chromatogram quality based on peak purity. The detected peaks were then identified by comparing the mass spectra of each chromatogram with the Mass Spectral Library (National Institute of Standards and Technology: NIST 20). The compounds were verified by measuring and comparing the Kovats retention indices based on the retention times of a saturated alkane mixture carried along with the samples (C9-C25; Sigma-Aldrich) as reported by (Guo et al., 2019, 2020). Peak areas were normalized according to the fresh weight of each analyzed sample. Six dilutions of the external standards – sabinene, α-pinene, linalool, methylsalicylate, β-caryophyllene, α-humulene, geraniol, and bornyl acetate were used for the quantification of the compounds.

### Statistical analyses

The data were logarithmically transformed for statistical analysis to fulfill assumptions of the normal distribution. Before the log transformation the data was tested for the normal distribution by using Shapiro-Wilk test. To visualize the terpenoid content of all analyzed tansy tissues, we plotted a heatmap with a “Euclidean” distance measure between different plants tissues, using the normalized concentration of mono- and sesquiterpenoids separately. Each colored cell in the heatmap corresponds to a compound concentration, e.g. compounds in the dark red cell on the map have the highest abundance in a sample. The partial least squares discriminant analysis (PLS-DA) model of multivariate statistics was used as described in Bertić et al. (2021; 2023). In our study, the model helped us to find tissue-specific discriminant mono- and sesquiterpenoid compounds and to show the separation between tissues. The PLS-DA model was calculated by maximizing the covariance between the X and Y variables, X variables (32 compounds of the monoterpenoid group and 41 compounds of the sesquiterpenoid group) and Y variables (leaflets, midrip, rhizome, coarse root and fine root tissues). The model was plotted separately for monoterpenoid and sesquiterpenoid variables as shown in Figure 1b and 2b. Each plot shows how the modelled observations lie in the X-space. Observations that are close together are more similar than observations that are relatively far apart. Components 1 and 2 explain the largest and second largest variation in the X-space. The fitness of the model was cross validated by the R2 and Q2 values, which indicated how well the variation in a variable is explained and how well a variable can be predicted. Note that well-modelled variables have high levels (as a reference: 0.5 or above) of R2 and Q2 values. The result of the PLS-DA model fitness is given in the legend of each plot.

**Figure 1:**
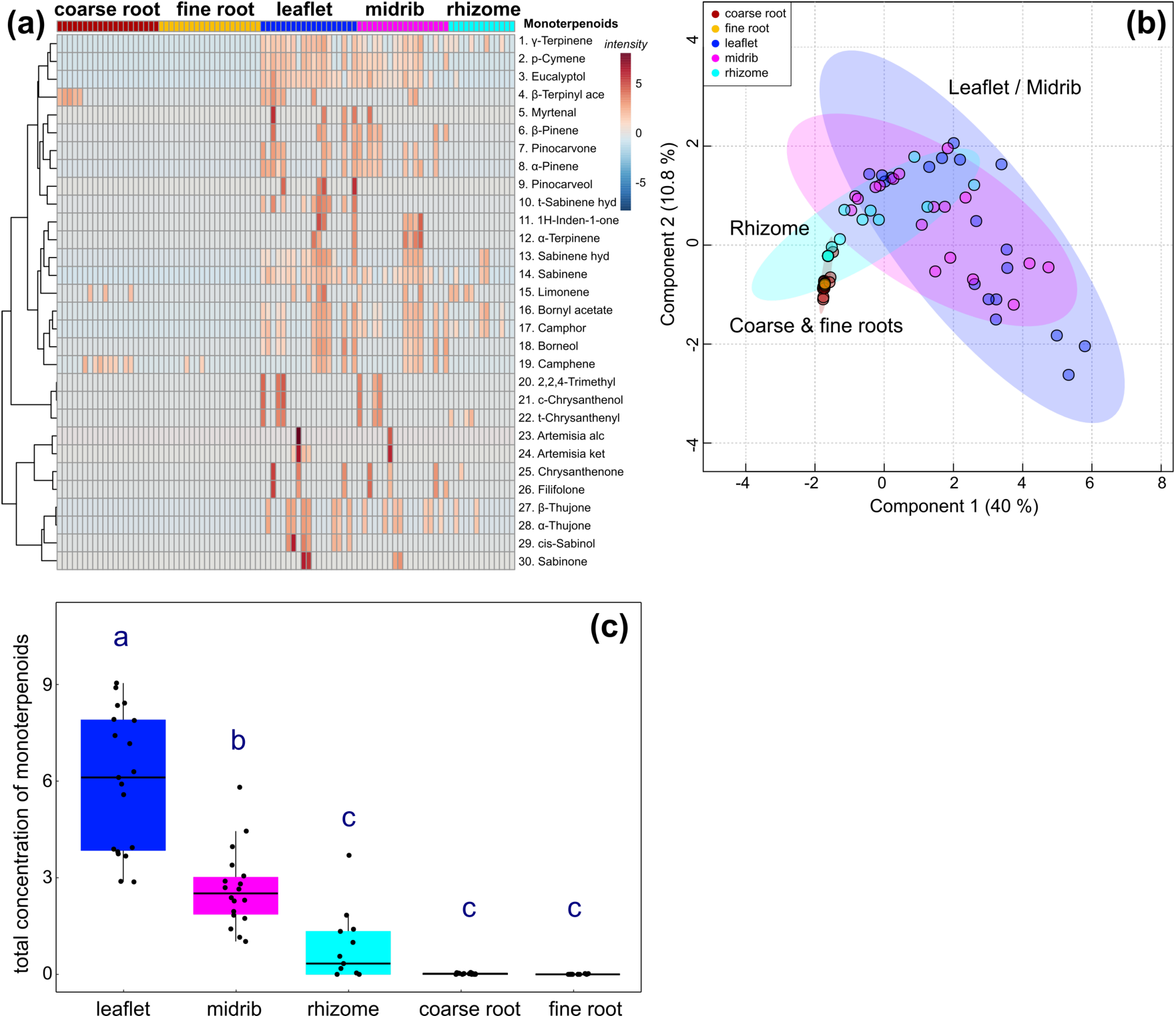
Tissue atlas of monoterpenoids presented in all tissues of untreated plants; **(a)** heatmap shows the monoterpenoid compounds across all the plants analyzed. The distribution of monoterpenoids are displayed among coarse root, fine root, leaf, midrib and rhizome tissues that are indicated in the legend accordingly. The intensity column on the right shows the concentration variation of the compounds. For instance, dark red cells on the map indicates the highest abundance and blue cells show that no compound is present. **(b)** PLS-DA exhibits a clustering between the tissues separated by the most discriminant mass features of monoterpenoids. The two components describe the data with total of 51.6% of explained variance. PLS model fitness: *R*^2^*X*(cum) = 0.821, *R*^2^*Y*(cum) = 0.556, *Q*^2^(cum) = 0.374 **(c)** Total concentration of monoterpenoids is significantly (one-factorial ANOVA, p < 0.05) elevated between leaflets, midrib, and rhizome, but not in coarse and fine root tissues. Abundance of monoterpenoids is very low in coarse and fine roots. Tissues labelled with the same letter do not differ significantly at p < 0.05 (posthoc test).

**Figure 2:**
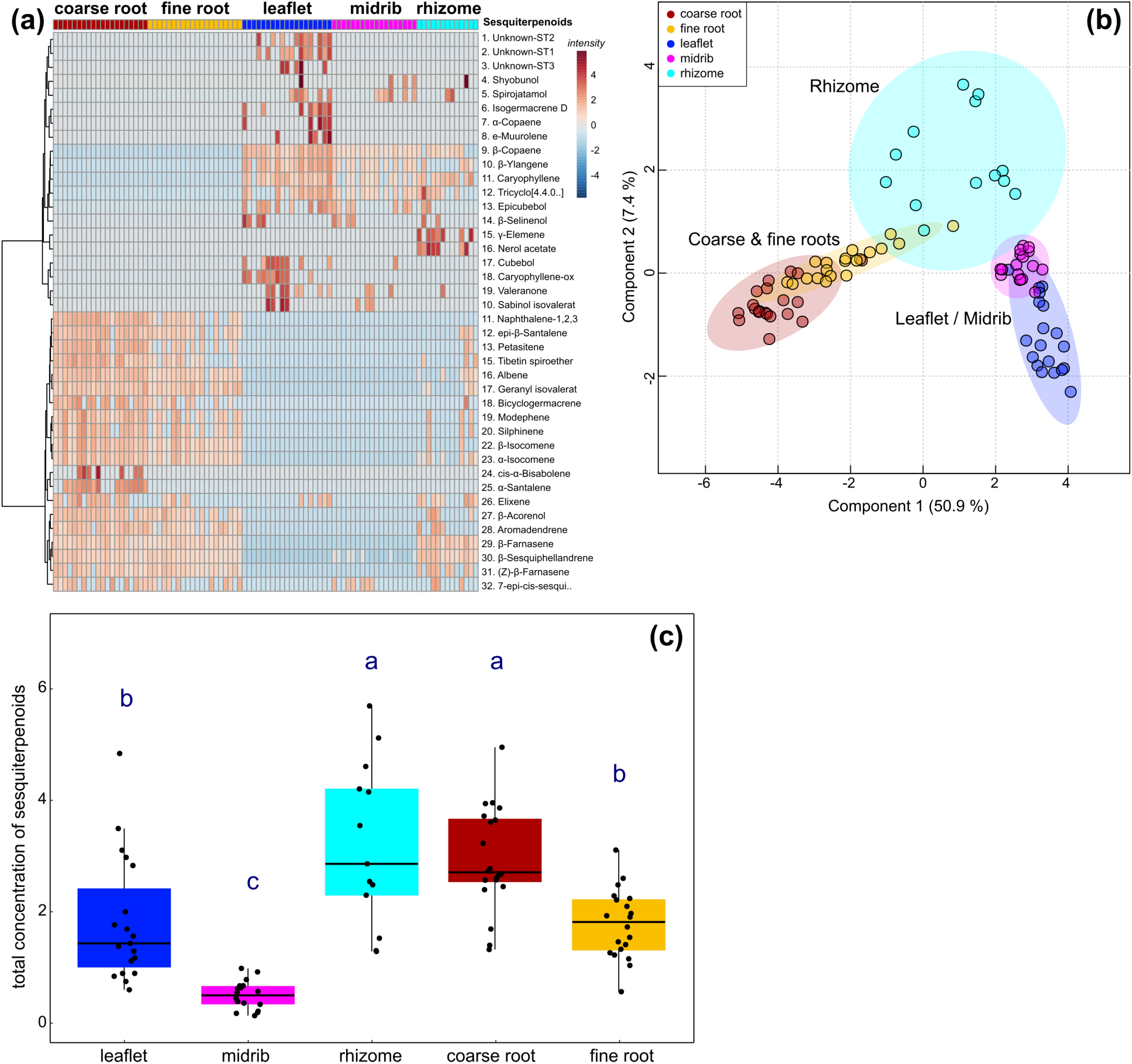
Tissue atlas of sesquiterpenoids identified in all tissues of untreated plants; **(a)** heatmap displays the distinct sesquiterpenoid profile among all the plants and tissues analyzed. The intensity column on the right describes the abundance of compounds in each sample **(b)** PLS-DA shows a clear separation between the tissues according to their sesquiterpenoid content. The two components show the data with total of 60.3% of explained variance. PLS model fitness: *R*^2^*X*(cum) = 0.720, *R*^2^*Y*(cum) = 0.740, *Q*^2^(cum) = 0.709 **(c)** total concentration of sesquiterpenoids were tested (one-factorial ANOVA, p < 0.05) between leaf, midrib, rhizome, coarse and, fine root tissues. Tissues labelled with the same letter do not differ significantly at p < 0.05 (posthoc test).

Alluvial diagram was plotted to observe the terpenoids flow from the shoots to the rhizomes and roots. On the X axes, three states of categorical variable, shoot, rhizome and root zones are presented. The flow of terpenoid groups is presented in different colors and spans through the zones. Alluvial diagrams are majorly used for qualitative analyses to graphically display the associations between categorical variables.

Total concentration of mono- and sesquiterpenoids between five different tissues were tested by using posthoc test (p < 0.05). T-test was used to test total concentration of mono- and sesquiterpenoids between pip treated and untreated plants of each tissue. The induction effect of each individual compound between untreated and pip treated groups were tested by using one-way ANOVA (p < 0.05). Furthermore, the control group plants were pooled and clustered into monoterpenoid or sesquiterpenoid chemotypes using the “factoextra” package (Kassambara & Mundt, 2020a), “hclust” function with the “ward.D2” method of correlation distance in R as described by Rahimova & Neuhaus-Harr et al. (2024).

Statistical tests were carried out using R (version 4.3.2; R Core Team 2023) and MetaboAnalyst 6.0 (Pang et al., 2021). All graphs were produced using the R package ggplot2 (Wickham, 2016) and the resulting images were edited using the “Inkscape” (version 1.1.1) image processing software for enhanced resolution.

## Results

### Monoterpenoid compositions of different tissues in tansy

First, the terpenoid composition of untreated tansy plants was investigated. The heat map showing the monoterpenoid profile of leaflets, leaf midribs, rhizome, coarse and fine root tissues showed that monoterpenoids were predominant in leaf and midrib tissues. (Figure 1a). Most of the dominant compounds such as β-thujone, trans-chrysanthenyl acetate, camphor and some minor monoterpenoids were, however, also identified in the rhizome tissue. For example, β-thujone was dominant in the leaflets, but also in the midrib and rhizome of the same individuals, although the concentration varied between tissues. The total concentration of monoterpenoids was significantly (Tukey’s HSD, p < 0.05) higher in leaves than in midribs and rhizomes (Figure 1c). In contrast to shoots, roots showed lower levels of monoterpenoids, and only three compounds, camphene, limonene and α-terpinyl acetate were detected in coarse and fine roots in trace amounts. The differences between the monoterpenoid profiles of the different tissues were clearly separated in a PLS-DA (Figure 1b). Using cluster analysis, the plants of the tansy control group could be separated into four different classes of monoterpenoid chemotypes in leaflets and leaf midribs (Figure S1a). Class 1 was characterized by a mixture of two main compounds: 31% sabinene hydrate and 27% camphor. Class 2 was defined predominantly by 70% trans-chrysanthenyl acetate, class 3 by 75% β-thujone, and class 4 comprised a group of plants with a mixture of monoterpenoids (Figure S1b). This marked variation between monoterpenoid classes is consistent with our previous study (Rahimova et al., 2024), showing that intraspecific variation in terpenoids can be used to determine tansy chemotypes.

### Sesquiterpenoid compositions of different tissues in tansy

The sesquiterpenoid profiles of the shoot and root systems were clearly different, as shown in the heat map (Figure 2a). In the shoot system, leaflets and midribs contained a very similar sesquiterpenoid profile. Compounds such as β-copaene, β-caryophyllene, β-ylangene, epicubebol, and oxyganated sesquiterpenoids such as spirojatamol, β-selinenol and sabinol isovalerate were identified compounds in almost all individuals in leaf and midrib tissues. In contrast to the shoot sesquiterpenoid composition, root systems produced different groups of sesquiterpenoid compounds, such as β-sesquiphellandrene, β-farnesene, α-isocomene and β-isocomene in coarse and fine root tissues, albeit at different concentrations. The terpenoid pattern and content of the rhizomes was comparable to that of the shoot and root sesquiterpenoid mixture, but there were also unique compounds, i.e.nerol acetate and γ-elemene (Figure 2a), which were only present in the rhizomes. Shoot-specific compounds such as β-copaene, β-caryophyllene and β-ylangene, and root-specific compounds such as β-sesquiphellandrene, β-farnesene and α-isocomene were found in the rhizome profile. PLS-DA showed the clear separation of the sesquiterpenoid content among the analyzed tissues (Figure 2b). The total concentration of sesquiterpenoids was significantly lower for midribs compared to rhizomes, leaflets and coarse and fine roots (Tukey’s HSD, p < 0.05) (Figure 2c).

The leaf and midrib samples were categorized into four aboveground sesquiterpenoid chemotype classes. Each class is shown in Figure S2b in a different color. β-Copaene was the dominant compound in each class. Classes 2 and 3 contained unique compounds: 27% sabinolisovalerate and 13% valeranone in class 2 and 12% β-selinenol in class 3, along with the predominant β-copaene (Figure S2b). Sesquiterpenoids present in coarse and fine root tissues could also be grouped into four belowground chemotype classes (Figure S2c). β-Sesquiphellandrene was the most prevalent sesquiterpenoid in each class of belowground chemotype, followed by β-farnesene, geranylisovalerate and β-isocomene. However, the abundance of these compounds varied between classes (Figure S2c). Our previous study (Rahimova et al., 2024) demonstrated that sesquiterpenoid chemotypes defined for leaves (and now also in the roots) do not exhibit pronounced variation in content, unlike the monoterpenoid chemotypes. Instead, a single dominant sesquiterpenoid is detected and slightly varied in the abundance across all individuals. We also plotted a tanglegram diagram (Fig. S2a) to visualize the data and investigate links between the aboveground (AG) and belowground (BG) sesquiterpenoid chemotype classes. Our analysis revealed no direct link between AG and BG chemotype classes, likely due to the sesquiterpenoids being synthesized by different terpene synthases in shoot and root tissues.

### Metabolic atlas of terpenoids

In addition, we examined the changes in terpenoid profiles from the shoots to the rhizomes and roots and presented them in an alluvial plot (Figure 3). As shown in the diagram, the proportion of monoterpenoids (MT) and oxygenated monoterpenoids (oMT) was predominant in the aerial parts of the shoot, but decreased towards the rhizome and almost disappeared in the root zone (ND - not detected). The proportion and number of sesquiterpenoids (SQT) increased from the leaves towards the rhizome, while the number of oxygenated sesquiterpenoids (oSQT) decreased in the opposite direction. Roots showed a markedly different pattern of sesquiterpenoids from the aerial parts of the plant. The rhizomes, which act as a transition zone, contained a mixture of above- and below-ground sesquiterpenoids. This pattern of terpenoids, extending from the shoot through the rhizome to the root zone, provided compelling evidence that there are terpenoid gradients in tansy plants.

**Figure 3:**
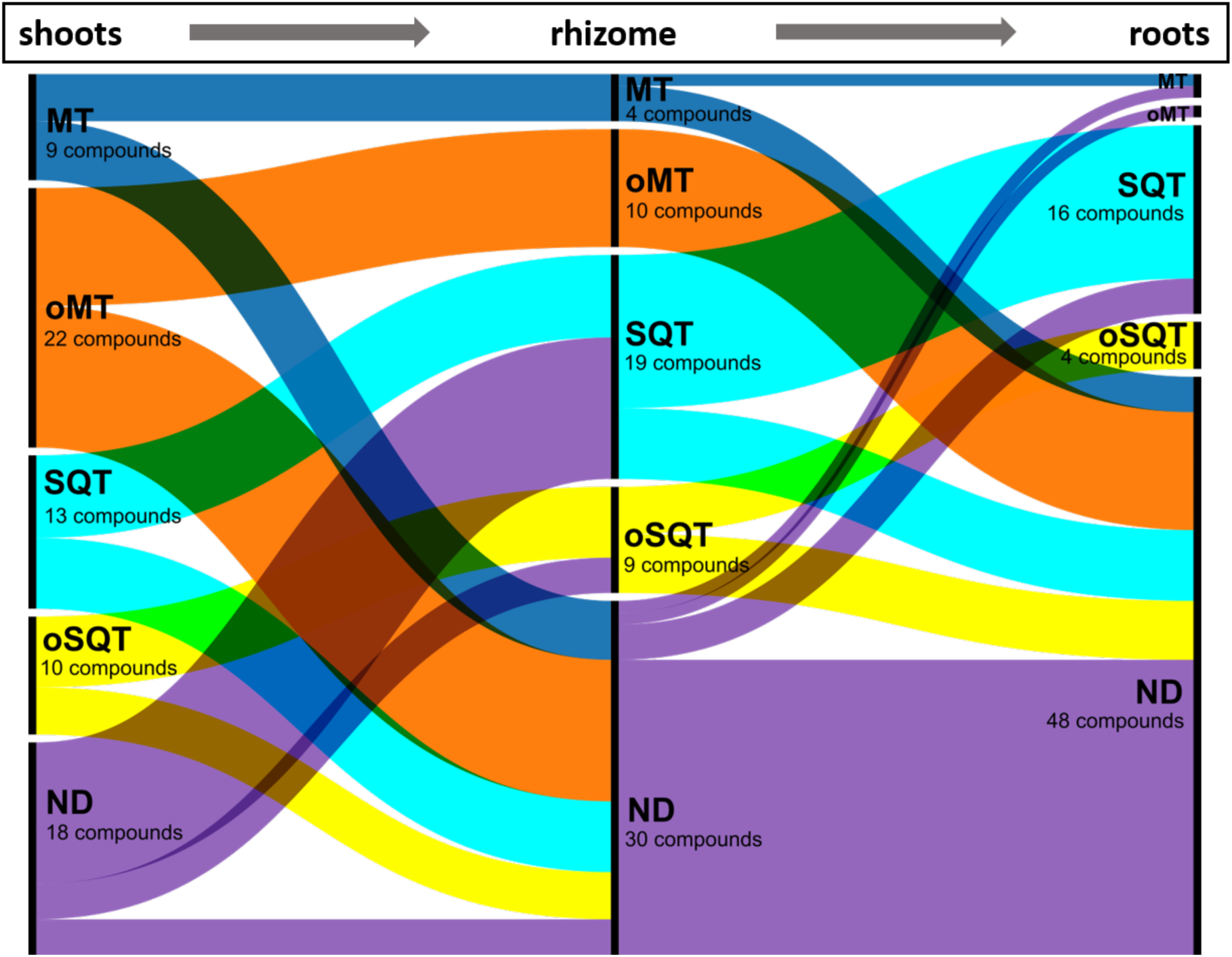
Alluvial plot shows the flow of terpenoids from shoots to rhizome and roots. The data are read from left to right as indicated by the arrows between the variables (shoots, rhizome and roots). Each categorical variable is shown in a different colour: MT - monoterpenoids in blue, oMT - oxygenated monoterpenoids in orange, SQT - sesquiterpenoids in green, oSQT - oxygenated sesquiterpenoids in red, ND - not detected in purple.

### Exogenously applied Pipecolic acid increased the induction of terpenoids

Following application of Pip to the plants, we observed an increase in the total concentration of mono- and sesquiterpenoids within three days of SAR stimulation in all tissues except rhizomes. However, t-tests demonstrated that the total concentration of monoterpenoids was significantly elevated by the application of Pip solely in the leaflets (F_x,y_, p = 0.006), but not in the midribs, rhizomes, coarse and fine roots (Figure 4a). In these tissues, only a tendency towards higher monoterpenoid contents could be observed. In contrast to the monoterpenoids, the sesquiterpenoids demonstrated a significant induction effect in almost all tissues, including leaflets, midribs, coarse roots and fine roots (t-tests, p < 0.001), with the exception of rhizomes (Figure 4b). Upon evaluation of each terpenoid in each tissue, no discernible difference was observed between the levels of terpenoids in the control group and the Pip-treated plant group. Of the identified monoterpenoids, only *π*-cymene exhibited a significant induction in the leaves following Pip application (one-way ANOVA, p = 0.05) (Figure 5a). Notably, no novel sesquiterpenoids not present in the control plants were induced by treatment with Pip.

**Figure 4:**
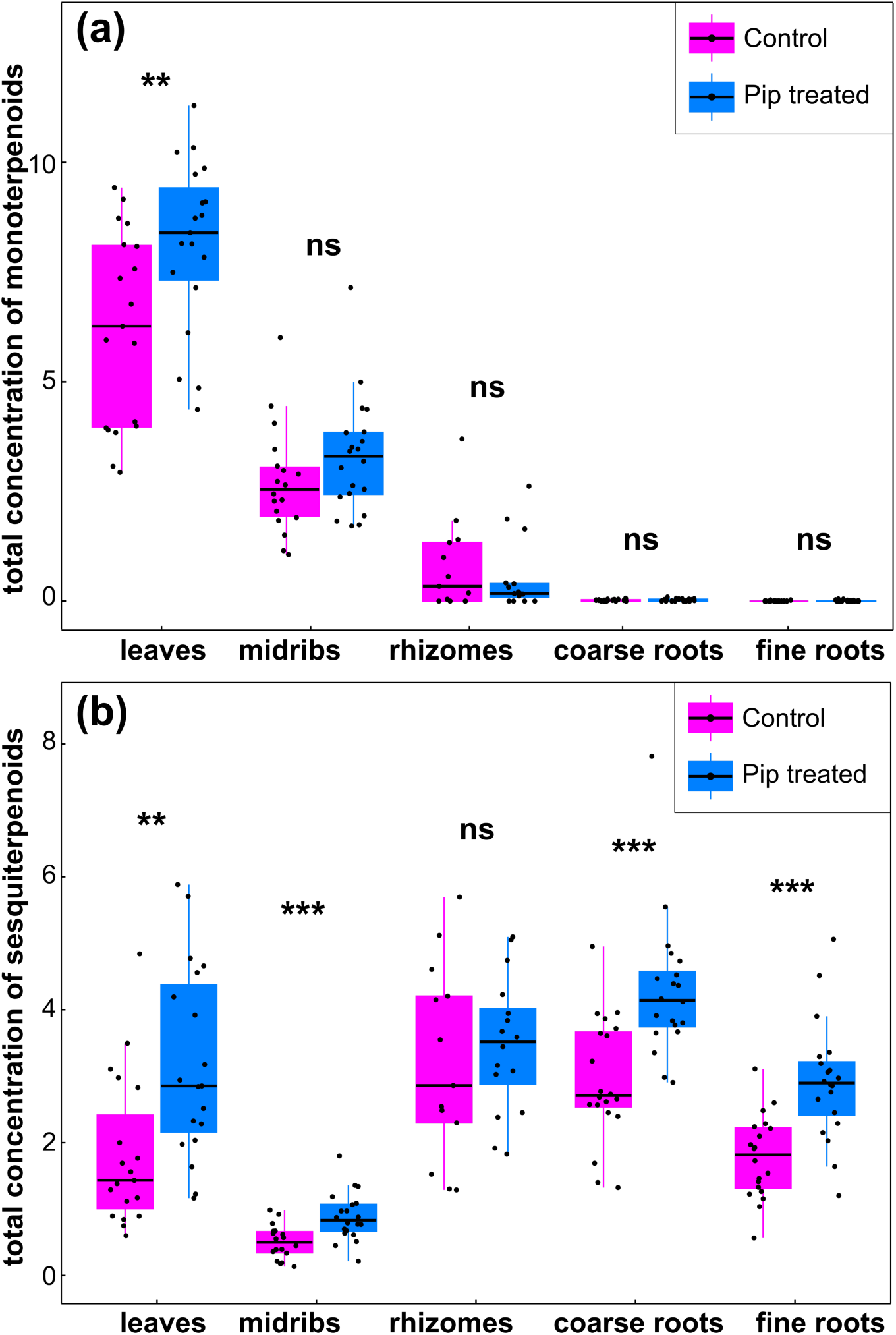
**(a)** Total concentration of monoterpenoids is significantly higher between control and Pip-treated plants for leaflets (t-test, p = 0.006), but not for midrib, rhizome, coarse root and fine roots. **(b)** Total concentration of sesquiterpenoids is significantly higher for all the tissues except for rhizomes (ns).

**Figure 5:**
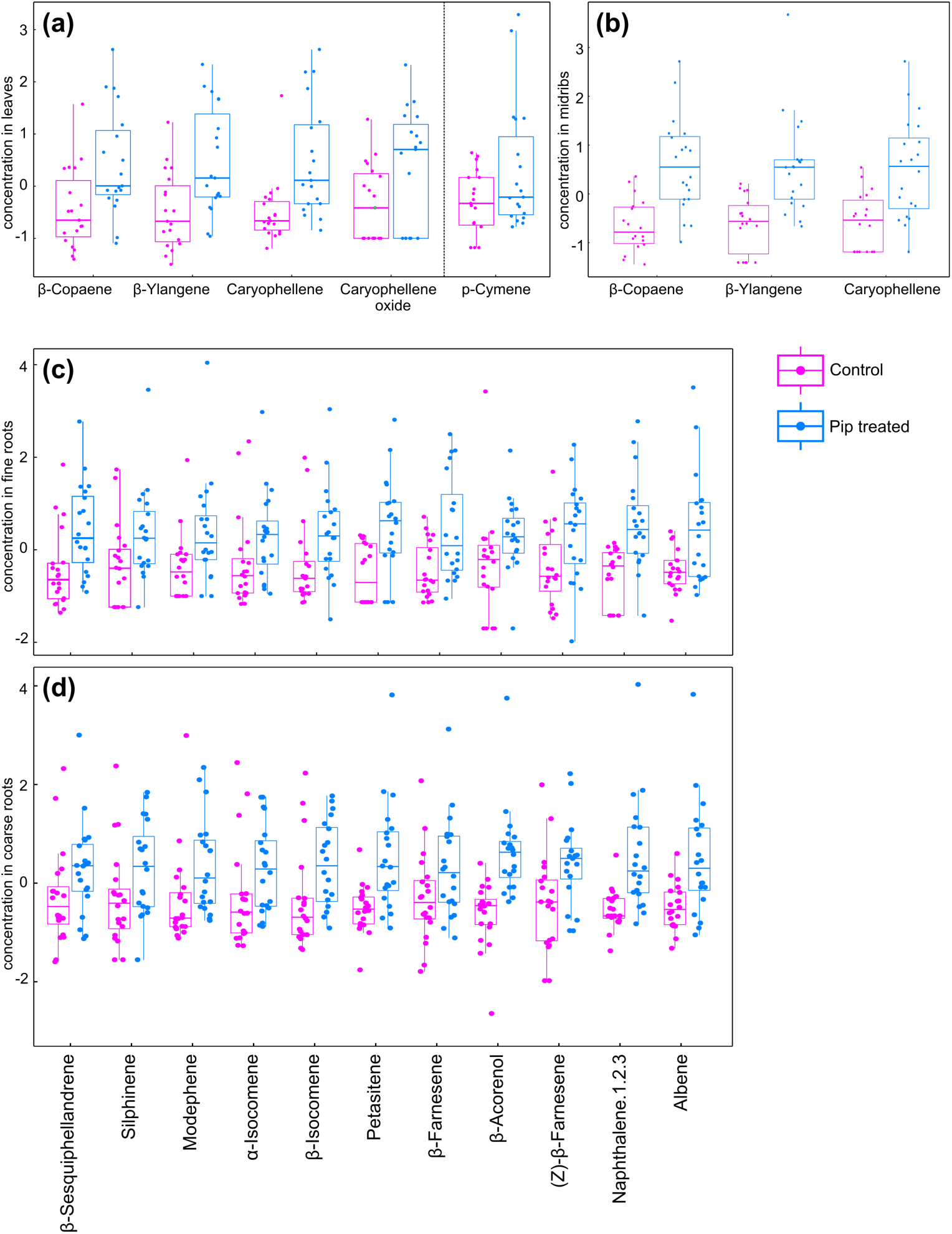
Concentration of significantly increased terpenoids between control (green boxes) and Pip-treated plants (red boxes). **(a)** Four sesquiterpenoid compounds, β-copaene, β-ylangene, caryophyllene, β-caryophyllene oxide (one-way ANOVA, p < 0.05) and one monoterpenoid compound, *π*-cymene (p = 0.05) in leaflets and **(b)** Three sesquiterpenoid compounds, β-copaene, β-ylangene, β-caryophyllene in midribs are strongly elevated. **(c)** 12 sesquiterpenoid compounds in fine roots and **(d)** 15 sesquiterpenoid compounds in coarse roots are significantly induced in Pip-treated plants. Y-axis (concentration) is log-scaled.

In contrast to the monoterpenoids, numerous individual sesquiterpenoid compounds exhibited a higher concentration in the shoot and root tissues of the Pip-treated plants. The sesquiterpenoid β-copaene was found to be the most abundant in the shoot tissues, with concentrations almost twice as high in the leaflets (one-way ANOVA, p = 0.004, Figure 5a) and midribs (one-way ANOVA, p < 0.001, Figure 5b) of the Pip-treated plants compared to those of the control group three days after the start of treatment. Furthermore, some low-abundance sesquiterpenoids, including β-ylangene, β-caryophyllene, and others, were observed to be significantly increased in the shoot tissues of Pip-treated plants after three days (Figure 5a,b, Table S1). It should be noted that the sesquiterpenoid compounds mentioned above are not present in the root tissues, therefore these compounds could not be tested statistically. The treatment with Pip led to a significant increase in the sesquiterpenoid contents of the root tissues compared to the shoot tissues and the rhizome, in which the concentrations only tended to increase. In addition to the dominant root sesquiterpenoids β-sesquiphellandrene and β-farnesene, 15 other individual sesquiterpenoid compounds were induced by Pip in the coarse and fine roots of the plants (Figure 5c,d, Table S1). The results of the statistical tests for each individual compound between Pip-treated and untreated plants are shown in Table S1.

## Discussion

The objective of this study was to investigate the effects of an exogenous Pip application on tansy, which mimics an aphid infestation (Agrawal, 1998, Kloth et al., 2016). Additionally, the study aimed to create a metabolic atlas of terpenoid distribution and chemical diversity by examining the terpenoid profiles in five different tissues: leaflets, leaf midribs, rhizome, coarse root and fine root, respectively. The distribution of terpenoids between tissues and their concentrations were found to vary considerably, confirming our initial hypothesis that there are tissue-specific differences in the overall terpenoid composition of tansy. Contrary to our expectations, we found that the application of Pip induced only low levels of monoterpenoids in aboveground plant organs, particularly in leaflets, within three days of treatment initiation. In contrast, sesquiterpenoids were strongly induced, with the greatest induction observed in belowground plant organs. This finding supports our second hypothesis that sesquiterpenoids are inducible components of orchestrated induced plant defenses. Our data offer some valuable insights into the inducibility of terpenoid mixtures in tansy in different tissues. They also provide some evidence for the involvement of different terpenoid groups in different ecological interactions involving these tissues in above- and below-ground.

Our analysis demonstrates the already known high intraspecific chemical diversity of terpenoids in the aboveground tissues of tansy plants. Furthermore, it shows that individuals can be categorized into chemotypic classes based on their patterns. This is in accordance with previous studies by Clancy et al. (2016), Rahimova et al. (2024) and Neuhaus-Harr et al. (2024). However, the roots of the tansy plant also exhibit a high degree of intraspecific diversity in their terpenoid profiles between different tissues. While the above-ground tissues are generally high in monoterpenoids, the below-ground tissues tend to be low in monoterpenoids (Kleine & Müller, 2013; Muchlinski et al., 2019). This is not an uncommon occurrence and is consistent with other studies demonstrating that monoterpenoids are produced in roots in a highly cell-type-specific manner (Chen et al., 2011). Despite their low abundance, monoterpenoids play an important role in root development and the interaction of roots with the rhizosphere (Lin et al., 2007). In contrast to monoterpenoids, the distribution of sequiterpenoids across the tissues is markedly different. They are less present in the above-ground tissues. Furthermore, their individual patterns differ less between plants, and the classification of plants into sesquiterpenoid chemotypes is less pronounced than for monoterpenoids. This observation is corroborated by a recent study of tansy chemotypes from across Germany (Rahimova et al., 2024). It also demonstrates that monoterpenoid and sequiterpenoid chemotypes are formed independently of each other in tansy leaves.

The metabolic atlas also indicates that distinct patterns of sequiterpenoids are formed in above-ground and below-ground tissues. In particular, sesquiterpenoids are only detectable in roots or green tissue. The data obtained in this study corroborate the findings of Kleine & Müller (2013), which also demonstrate the clear separation of mono- and sesquiterpenoids in tansy leaves and roots. The sequiterpenoids identified in their study can also be detected in our study. Furthermore, additional sequiterpenoids such as silphinene, modephene, β-Isocomene and many others were detected in the present study, which is likely due to the larger number of plants examined.

The differences in the terpenoid profiles of tansy roots and shoots are likely due to the different abiotic and biotic pressures on specialized metabolites in aboveground and belowground plant compartments (Kleine & Müller, 2013; van Dam, 2009). The highly volatile monoterpenoids, which are typically stored in glands and glandular trichomes in aboveground tissues, evaporate rapidly. In contrast, sesquiterpenoids (and other semi-volatile compounds, e.g. borneol) have longer C chains, higher boiling points and low Henry’s law constants (Mofikoya et al., 2019), making them less volatile and less soluble in water. Thus, sesquiterpenoids are not volatile enough to be emitted from undisturbed tissues and are more likely to be concentrated in the rhizosphere, whereas highly hydrophilic substances such as monoterpenoids would leach out more rapidly and dissipate into the soil (Qualls, 2005). This explains why monoterpenoids are abundant aboveground, where they can be used to communicate with aboveground organisms, including herbivores and their natural enemies or pollinators (Neuhaus-Harr et al., 2024; Sasidharan et al., 2023; Ziaja & Müller, 2023).

Distinct profiles of secondary metabolites have been observed in roots and shoots of other plant species. For instance, in *Brassica oleracea* L., the quantity of glucosinolate metabolites differs between the shoot and root, with the concentration of these compounds being higher in the roots than in the shoot (Kabouw et al., 2010). Such variations can also be demonstrated for phenolic compounds. A study on *Oenothera biennis* L. revealed that shoot tissue contains chlorogenic acid and flavonoids that were not observed in the roots, while root tissue contains ellagic acids that are not found in the shoots (Parker et al., 2012). Ristok and colleagues (2019) demonstrated that the metabolite content of the meadow herb species *Centaurea jacea* L, *Knautia arvensis*, (L.) Coult., *Leucanthemum vulgare* Lam. and *Plantago lanceolata* L. exhibited significant differences between shoots and roots. The chemical diversity of secondary metabolites was found in these examples to be higher in shoots than in roots.

It has been demonstrated that aphid feeding on plants stimulates the production of salicylic acid (SA) (Voelckel et al., 2004; Kloth et al., 2016), whereas leaf-feeding herbivores activate JA/ethylene metabolism (Walling, 2000). Therefore, we used pipecolic acid (Pip), a positive regulator for SA production and defense amplification (Navarova et al., 2013; Zeier, 2013) to demonstrate the impact of Pip production on the induction of systemic induced resistance (SAR) (Chen et al., 2018; Hartmann et al., 2018; Vlot et al., 2021) by examining the changes in the terpenoid pattern in the different tissues of tansy. This was conducted with the understanding that the initiation of the systemic defense reaction is clearly defined in response to this type of treatment, whereas the start of the reaction in response to natural aphid colonization is difficult to define, making the interpretation of pattern changes challenging (see Clancy et al., 2018).

The application of Pip led to an increase in subgroup-specific and tissue-specific increases in terpenoid levels. By mimicking biotic stress through the introduction of aphids, we demonstrated that terpenoids, particularly sesquiterpenoids, exhibited a significant increase in all plant tissues, with the greatest increase observed in root tissue in response to Pip. Pipecolic acid has been identified as a key triggering molecule for the induction of SAR in plants through the activation of defense responses (Bernsdorff et al., 2016; Hartmann et al., 2018). In many plant species, SAR induction results in the emission of stress-induced VOCs, which play a pivotal role in plant-to-plant communication, defense against herbivores and more generally in interaction with the environment (Pierik et al., 2014; Vlot et al., 2021). In *Arabidopsis*, monoterpenes (α/β-pinene and camphene) are increasingly emitted after SAR induction (Riedlmeier et al., 2017). In tomato (*Solanum lycopericum* L.) plants, it has been demonstrated that plant defense against infection with the Phytoplasma Potato Purple Top (PPT) is more effective after the application of the phytohormone SA (Wu et al., 2012). Another study demonstrated that exogenously applied Pip induces SAR in cucumber (*Cucumis sativus* L.) plants and enhances the activity of defense-associated enzymes against subsequent pathogen infection (Pazarlar et al., 2021).

Although the specific mechanisms of SAR induction in tansy have not yet been investigated and further studies are needed, especially with regard to genetic regulation by specific terpene synthases (TPS) and other backbone modifying enzymes, it is highly suggestive that the increasing concentration of terpenoids in leaflets, midribs and coarse and fine roots reflects an indication of an inducible defense response. However, this interpretation is complicated by the compartmentalization of terpenoids in tansy. Tansy leaves, including the midribs (the main target site of aphid attack), are characterized by a high density of glandular cells, which serve as storage sites for monoterpenoids in particular (Guerreiro et al., 2016; Bergman et al., 2023). Jakobs et al. (2018) showed plant organ-specific responses to aphid infestation, for example, the composition of sugars and organic acids in the phloem exudate of tansy was particularly influenced by aphids compared to leaves. This shows that metabolic differences in different tansy organs play an important role in tansy-insect interaction.

The presence of constitutively present monoterpenoid reservoirs, which can be used as stable chemical markers for the differentiation of chemotypes (see here, as well as Clancy et al. 2016, Rahimova et al., 2024), makes it challenging to demonstrate the induction of stress-induced monoterpenes. The available data indicate a weak tendency for an increase in monoterpenoid content following the application of the Pip, which suggests that the formation of new monoterpenes was relatively low compared to the existing pool.

In contrast, the localization of sesquiterpenoids in glandular cells is less well documented than that of monoterpenoids. However, previous comparative analyses of hexane extracts from tansy leaves with VOC emission measurements (Clancy et al., 2016, 2020) indicate that these compounds are increasingly synthesized under herbivore stress. The cellular localization of mono- and sesquiterpenoids in the rhizomes and roots remains unknown. No storage tissues or subcellular structures with terpenoids (such as vesicles) have been described in either tissue. Due to their high lipophilicity, terpenoids could be dissolved in biomembranes and extracted by hexane (Huang et al., 2008; Clancy et al., 2016). The results presented here clearly demonstrate the induction of this terpenoid subgroup in tansy, with a particularly strong response observed in the roots.

Terpene synthases (TPS) are, together with backbone modifying enzymes downstream, responsible for the great diversity of mono- and sesquiterpenoids. Terpene synthases form a group of enzymes that catalyze the conversion of terpenoid compounds from isoprenoid precursors (Chen et al., 2011; Degenhardt et al., 2009). The TPS gene family has been identified in various plant species, including the leaf tissues of the tansy, which is characterized by mono- and sesquiterpenoid synthases (Clancy et al., 2020). It can be assumed that different TPS genes, which regulate organ-specific biosynthesis of terpenoids, are responsible for the observed differences in tissue-specific terpenoid patterns as well as for the inducibility by Pip. For instance, in maize plants, various TPS enzymes are expressed exclusively in shoot and root tissue (Köllner et al., 2009). This work further demonstrates that these TPS enzymes produce sesquiterpenoids that are consistently induced by herbivores. As evidenced by the example of β-caryophyllene, these sesquiterpenoids play a role in the defense against the western corn rootworm by attracting insect-killing nematodes (Degenhard et al., 2009). Another study has demonstrated that chamomile plants (*Matricaria chamonmilla* L.) produce the common terpenoid hydrocarbons (β-caryophyllene, germacrene A, germacrene D and β-ocimene) in aerial organs. Furthermore, these compounds differ from several tricyclic hydrocarbons that are exclusively synthesized in roots, such as the sesquiterpenoid α-isocimene (Irmisch et al., 2012). Another example is provided by black mustard (*Rhamphospermum nigrum* L. AlShehbaz). Here, root herbivory leads to the formation of β-farnesene (Soler et al., 2007). Only recently, Lackus et al. (2021) detected a TPS gene (*PtTPS5*) that is induced in poplar roots during pathogen infection, which also indicates the involvement of sequiterpenoids in the pathogen defense of roots.

The current patterns of terpenoids in the different tissues of tansy and the different inducibility by simulated SAR encourage further investigation of these patterns using functional genomics. Expression analysis of TPS with GUS (β-glucuronidase) reporter-gene plants of *Artemisia annua* L., a species closely related to tansy, which also belongs to the Asteraceae, have provided information on the possible distribution of sesquiterpenoids in plant tissues (Wang et al., 2013). The promoter activity of β-caryophyllene synthase is observed at different stages of flower development, with the highest expression at full flowering. T-shaped trichomes, leaf primordia and leaf ribs of older leaves also show GUS staining, whereas no promoter activity was found in glandular cells. In roots, promoter activity was observed in vascular tissue. Other TPS, such as a α-farnesene synthase, are also expressed in roots, flowers and leaves, but also in glandular cells. Other TPS show different expression patterns and are absent in roots, for example. This shows that the metabolic atlas of terpenoid distribution in tansy presented here can probably be functionally explained by analyzing the very heterogeneous tissue-specific activity of the TPS. Since it is not clear where terpenoids might be localized in root and rhizome tissues, as no storage structures in roots have been reported so far, single cell metabolomics analyses could help to reveal the specialized cell types in tansy. For example, Li and colleagues (2024) showed that a cell type-specific transcription factor regulates the expression of two idioblast-specific biosynthetic genes in the monoterpenoid indole alkaloid (MIA) pathway of *Catharanthus roseus* L. (Madagascar periwinkle), providing insights into cell type-specific metabolic regulation. Single cell metabolomics of phloem tissues may be of particular importance, as phloem tissues have been shown to be preferred by aboveground feeding insects (Jacobs et al., 2018).

With the tansy genome available in draft form (personal communication from Andrea Bräutigam, University of Bielefeld, Germany), the necessary prerequisites for such a study have been created to understand the role of tissue-specific chemodiversity and its inducibility in the interaction of tansy with its communities. With such a study it will also be possible to functionally link terpenoid pattern to the role they play in the interaction with aphids, ants and arthropod metacommunities in general (Rahimova et al., 2024; Neuhaus-Harr et al. 2024; Ojeda-Prieto et al., 2024).

## Conclusions

The present study demonstrated that terpenoid profiles exhibit significant variation in compound content and concentration between different plant tissues, including leaflets, leaf midribs, rhizomes, coarse roots, and fine roots of tansy, respectively. These findings suggest that tansy shoot and root tissues may be subject to distinct challenges imposed by contrasting abiotic and biotic factors, including belowground feeding root herbivores, microorganisms, and aboveground insect populations. Furthermore, our findings indicate that pipecolic acid application mimicking SAR induction increases terpenoid levels in tansy tissues, suggesting enhanced defense responses that are likely regulated by specific enzymes. Our results highlight that the terpenoid diversity of tansy plants extends beyond intraspecific differences between individuals, but also occurs within individuals. Our study provides a foundation for further investigations into how the environment selects for chemical fingerprints in different plant tissues and, subsequently, how it mediates plant-associated communities.

## Acknowledgments

The authors wish to thank Ulrich Junghans and Ina Zimmer for their support during plant propagation and harvesting. This work was supported by grants of the Deutsche Forschungsgemeinschaft DFG to JPS (SCHN653/7-2) and WWW (WE3081/25-2 and WE3081/40-1).

## Contribution of authors

JPS conceived and designed the study. JPS, RH, and WWW reviewed, edited, and supervised the study. BW conducted the calibration and GC-MS analyses. HR performed the extraction, analyzed the data, and wrote the manuscript with substantial input from JPS, RH, and WWW.

## Supplementary Figures and Tables

**Figure S1:**
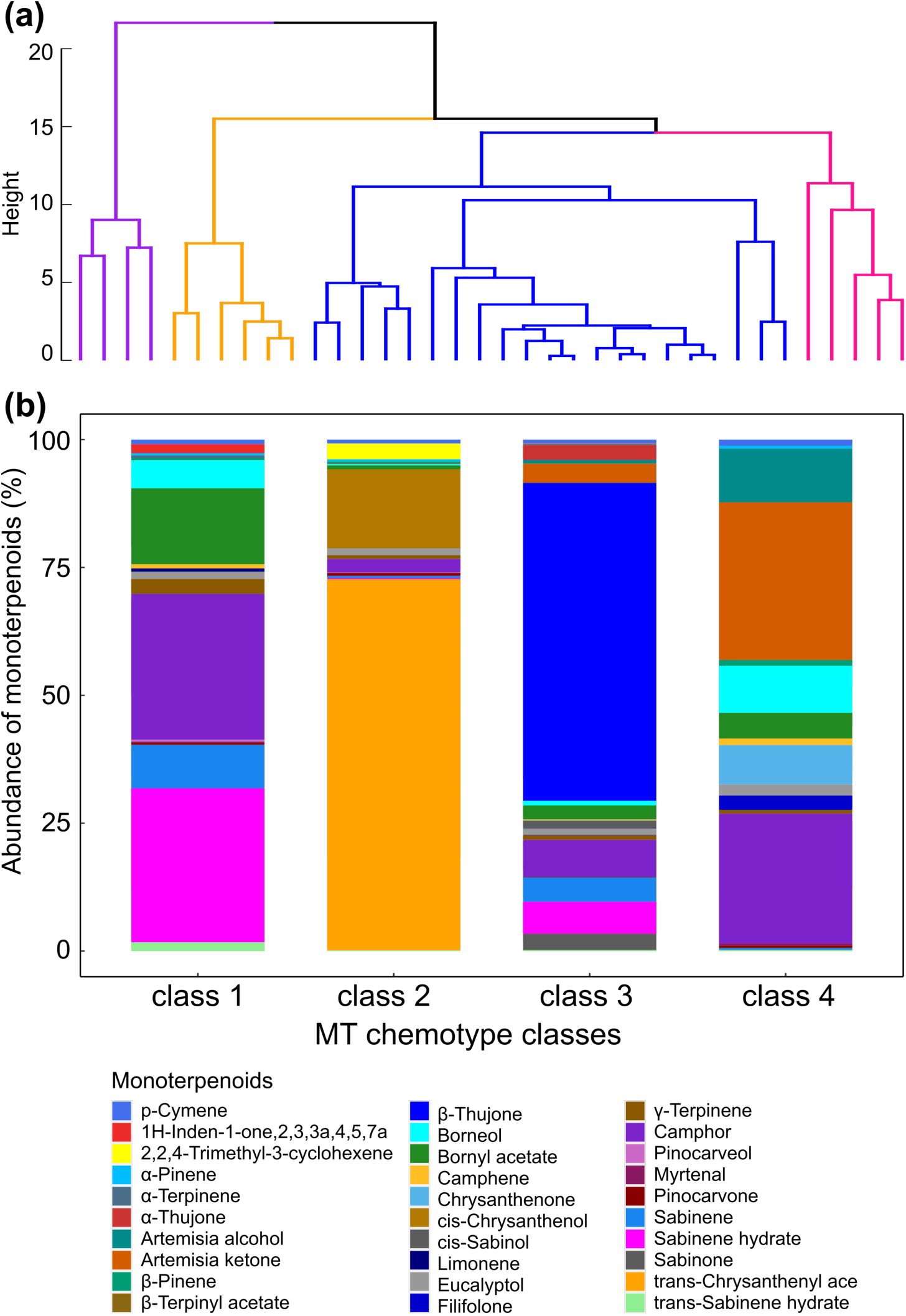
(a) Hierarchical cluster analysis of monoterpenoid compounds from leaf and midrib tissues of control plants. Four main classes are identified and each cluster is shown in a different colour; class 1 purple, class 2 orange, class 3 blue and class 4 magenta. Data are normalized to log scale. (**b**) The abundance of each representative compound of each class is presented in a stacked bar plot.

**Figure S2:**
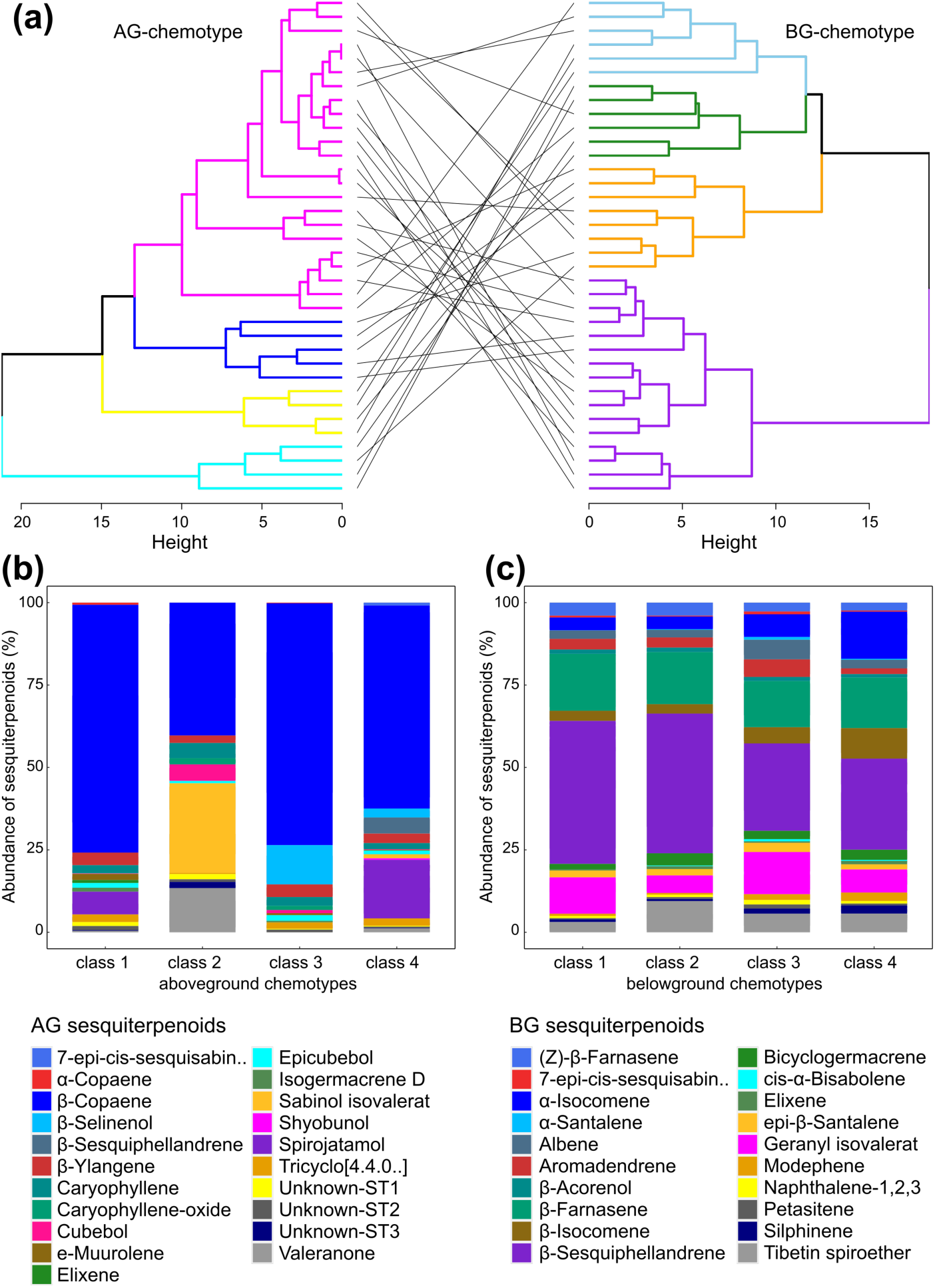
(**a**) Hierarchical cluster analysis shows that sesquiterpenoid compounds from leaf and midrib samples clustered into aboveground chemotypes (AG-chemotypes), shown in the left site tree with four different classes highlighted in different colours: From bottom to top, class 1 cyan, class 2 yellow, class 3 blue, class 4 magenta. Coarse and fine root samples clustered into belowground chemotypes (BG-chemotypes), shown on the right site tree in four colours: class 1 purple, class 2 orange, class 3 forest green, class 4 light blue. The tanglegram shows the relationships between the AG and BG chemotype classes. The abundance of sesquiterpenoids in AG (**b**) and BG (**c**) chemotypes is shown for each class in a stacked bar plot.

**Table S1:** Statistic (F-value) and p-value for individual terpenoid compounds tested between Pip-treated and untreated plants using a one-way ANOVA. Bold letters indicate significant p-values (p <0.050). Dashes indicate absence of the compound in the respective compartment. NA indicates that individuals with the mentioned compound are not enough for reliable statistical tests.

| Terpenoids | Leaf | Midrib | Rhizome | Coarse root | Fine root |
| --- | --- | --- | --- | --- | --- |
|  | F (p-value) | F (p-value) | F (p-value) | F (p-value) | F (p-value) |
| α-Pinene | 0.09 (0.77) | 0.04 (0.85) | - | - | - |
| Camphene | 0.63 (0.43) | 0.26 (0.61) | - | 0.17 (0.68) | 0.11 (0.75) |
| Sabinene | 1.47 (0.23) | 1.40 (0.24) | 1.65 (0.21) | - | - |
| β-Pinene | 0.70 (0.41) | 2.58 (0.12) | - | - | - |
| α-Terpinene | 0.33 (0.57) | 1.43 (0.24) | - | - | - |
| π-Cymene | **4.24 (0.05)** | 2.72 (0.11) | 1.58 (0.22) | - | - |
| Limonene | 0.23 (0.63) | 0.05 (0.82) | 1.52 (0.23) | 2.71 (0.11) | NA |
| β-Terpinyl acetate | 1.01 (0.32) | 0.004 (0.95) | - | NA | - |
| Eucalyptol | 0.67 (0.42) | 0.02 (0.89) | 0.26 (0.62) | - | - |
| Artemisia ketone | 0.40 (0.53) | 0.30 (0.58) | 0.81 (0.37) | - | - |
| γ-Terpinene | 0.58 (0.45) | 0.01 (0.92) | 0.97 (0.33) | - | - |
| Sabinene hydrate | 0.54 (0.47) | 0.30 (0.59) | 2.00 (0.17) | - | - |
| Artemisia alcohol | 0.34 (0.56) | 0.16 (0.69) | 0.81 (0.37) | - | - |
| Filifolone | 0.70 (0.41) | 0.67 (0.42) | - | - | - |
| trans-Sabinene hydrate | 0.02 (0.90) | 0.90 (0.35) | - | - | - |
| α-Thujone | 1.31 (0.26) | 1.62 (0.21) | 0.06 (0.81) | - | - |
| β-Thujone | 2.10 (0.16) | 2.30 (0.14) | 0.02 (0.88) | - | - |
| cis-Chrysanthanol | 1.60 (0.22) | 2.64 (0.11) | - | - | - |
| Chrysanthone | 0.84 (0.36) | 1.01 (0.31) | 0.28 (0.60) | - | - |
| 2,2,4-Trimethyl-3-cyclohexene-1-carbaldehyde | 1.32 (0.26) | 2.44 (0.13) | - | - | - |
| cis-Sabinol | 0.02 (0.88) | 1.90 (0.17) | - | - | - |
| L-Pinocarveol | 2.14 (0.15) | - | - | - | - |
| Camphor | 0.60 (0.46) | 0.62 (0.44) | 1.50 (0.23) | - | - |
| Sabinone | 0.001 (0.98) | 0 (0.1) | - | - | - |
| Pinocarvone | 1.19 (0.28) | 0.22 (0.64) | - | - | - |
| 1H-Inden-1-one, 2,3,3a,4,5,7a-hexahydro-5-methyl- | 2.50 (0.12) | 0.44 (0.51) | - | - | - |
| Borneol | 1.72 (0.20) | 1.12 (0.30) | - | - | - |
| D-Verbenone | 0.37 (0.54) | - | - | - | - |
| Myrtenal | 2.82 (0.10) | 0.50 (0.48) | - | - | - |
| trans-Chrysanthenyl acetate | 0.93 (0.34) | 1.67 (0.21) | 3.52 (0.07) | - | - |
| Bornyl acetate | 0.002 (0.97) | 0.02 (0.90) | 1.64 (0.21) | - | - |
| Elixene | **4.36 (0.04)** | - | 0.01 (0.93) | 0.31 (0.58) | 0.02 (0.90) |
| α-Copaene | **14.0 (<0.001)** | - | - | - | - |
| β-Ylangene | **11.1 (0.002)** | **19.5 (<0.001)** | NA | - | - |
| β-Caryophyllene | **12.0 (0.001)** | **15.6 (<0.001)** | NA | - | - |
| Tricyclo[4.4.0.0<sup>2,7</sup>]decane, 1-methyl-3-methylene | **4.75 (0.04)** | **5.84 (0.02)** | 0.56 (0.46) | - | - |
| Isogermacrene D | **11.1 (0.002)** | - | - | - | - |
| ε-Muurolene | **7.44 (0.01)** | 2.99 (0.09) | - | - | - |
| Unknown-ST1 | 3.29 (0.08) | - | - | - | - |
| Unknown-ST2 | 2.25 (0.14) | - | - | - | - |
| β-Copaene | **9.05 (0.005)** | **23.3 (<0.001)** | 3.12 (0.09) | - | - |
| Unknown-ST3 | 3.22 (0.08) | - | - | - | - |
| Bicyclogermacrene | 3.13 (0.09) | - | 0.80 (0.39) | 1.07 (0.31) | 2.20 (0.15) |
| Sabinol isovalerate | 1.45 (0.24) | 0.01 (0.90) | - | - | - |
| Epicubebol | 0.15 (0.70) | 3.31 (0.07) | 0.35 (0.56) | - | - |
| Caryophyllene oxide | **4.70 (0.04)** | - | - | - | - |
| Spirojatamol | 0.35 (0.55) | 0.03 (0.87) | 1.70 (0.20) | - | - |
| Caryophylladienol II | NA | - | - | - | - |
| Cubebol | 0.05 (0.82) | 1.02 (0.32) | - | - | - |
| β-Selinene | 0.003 (0.96) | 0.14 (0.71) | 0.38 (0.55) | - | - |
| Valeranone | 0.001 (0.97) | 0.11 (0.74) | 0.46 (0.50) | - | - |
| Shyobunol | 1.29 (0.26) | 0.02 (0.87) | 0.00 (0.99) | - | - |
| Aromandendrene | - | - | 1.10 (0.30) | **5.62 (0.02)** | 1.83 (0.18) |
| 7-epi-cis-Sesquisabinene hydrate | - | **5.07 (0.03)** | 0.14 (0.71) | 1.41 (0.24) | 0.01 (0.92) |
| β-Sesquiphellandrene | - | 0.20 (0.66) | 0.01 (0.93) | **Wilcoxon (0.02)** | **9.46 (0.004)** |
| Silphinene | - | - | 0.01 (0.94) | **4.02 (0.05)** | **5.81 (0.02)** |
| Modephene | - | - | 0.12 (0.73) | **5.55 (0.02)** | **6.58 (0.01)** |
| α-Isocomene | - | - | 0.01 (0.95) | **Wilcoxon (0.03)** | **Wilcoxon (0.01)** |
| β-Isocomene | - | - | 2.62 (0.12) | **6.53 (0.02)** | **5.66 (0.02)** |
| Petasitene | - | - | 2.19 (0.15) | **14.9 (<0.001)** | **14.8 (<0.001)** |
| β-Farnesene | - | - | 1.20 (0.28) | **4.02 (0.05)** | **10.6 (0.002)** |
| β-Acorenol | - | - | 0.09 (0.76) | **25.6 (<0.001)** | **4.13 (0.05)** |
| (Z)-β-Farnesene | - | - | 2.55 (0.12) | **7.82 (0.01)** | **6.85 (0.01)** |
| Tibetin spiroether | - | - | 0.01 (0.95) | **13.7 (0.001)** | **14.1 (<0.001)** |
| Naphthalene, 1,2,3,4,4a,7-hexahydro-1,6-dimethyl | - | - | - | **16.1 (<0.001)** | **19.8 (<0.001)** |
| Albene | - | - | 2.71 (0.11) | **12.1 (0.001)** | **10.3 (0.003)** |
| α-Santalene | - | - | - | **14.9 (<0.001)** | NA |
| Epi- β-Santalene | - | - | 0.82 (0.37) | **6.58 (0.01)** | 0.83 (0.37) |
| Geranyl isovalerate | - | - | 0.02 (0.88) | **4.37 (0.04)** | 3.28 (0.08) |
| cis-α-Bisabolene | - | - | - | **38.0 (<0.001)** | NA |
| γ-Elemene | - | - | 2.75 (0.11) | - | - |
| Nerol acetate | - | - | 0.11 (0.75) | - | - |

## Notes

### Competing Interest Statement

The authors have declared no competing interest.

